# Mosaic Plasmids are Abundant and Unevenly Distributed Across Prokaryotic Taxa

**DOI:** 10.1101/428219

**Authors:** Mitchell W. Pesesky, Rayna Tilley, David A. C. Beck

## Abstract

Mosaic plasmids, plasmids composed of genetic elements from distinct sources, are associated with the spread of antibiotic resistance genes. Transposons are considered the primary mechanism for mosaic plasmid formation, though other mechanisms have been observed in specific instances. The frequency with which mosaic plasmids have been described suggests they may play an important role in plasmid population dynamics. Our survey of the confirmed plasmid sequences available from complete and draft genomes in the RefSeq database shows that 46% of them fit a strict definition of mosaic. Mosaic plasmids are also not evenly distributed over the taxa represented in the database. Plasmids from some genera, including *Piscirickettsia* and *Yersinia*, are almost all mosaic, while plasmids from other genera, including *Borrelia*, are rarely mosaic. While some mosaic plasmids share identical regions with hundreds of others, the median mosaic plasmid only shares with 8 other plasmids.

When considering only plasmids from finished genomes (51.6% of the total), mosaic plasmids have significantly higher proportions of transposases and antibiotic resistance genes. Conversely, only 56.6% of mosaic fragments (DNA fragments shared between mosaic plasmids) contain a recognizable transposase, and only 1.2% of mosaic fragments are flanked by inverted repeats. Mosaic fragments associated with the IS26 transposase are 3.8-fold more abundant than any other sequence shared between mosaic plasmids in the database, though this is at least partly due to overrepresentation of *Enterobacteriaceae* plasmids.

Mosaic plasmids are a complicated trait of some plasmid populations, only partly explained by transposition. Though antibiotic resistance genes led to the identification of many mosaic plasmids, mosaic plasmids are a broad phenomenon encompassing many more traits than just antibiotic resistance. Further research will be required to determine the influence of ecology, host repair mechanisms, conjugation, and plasmid host range on the formation and influence of mosaic plasmids.

**Author Summary:** Plasmids are extrachromosomal genetic entities that are found in many prokaryotes. They serve as flexible storage for genes, and individual cells can make substantial changes to their characteristics by acquiring, losing, or modifying a plasmid. In some pathogenic bacteria, such as *Escherichia coli*, antibiotic resistance genes are known to spread primarily on plasmids. By analyzing a database of 8,592 plasmid sequences we determined that many of these plasmids have exchanged genes with each other, becoming mosaics of genes from different sources. We next separated these plasmids into groups based on the organism they were isolated from and found that different groups had different fractions of mosaic plasmids. This result was unexpected and suggests that the mechanisms and selective pressures causing mosaic plasmids do not occur evenly over all species. It also suggests that plasmids may provide different levels of potential variation to different species. This work uncovers a previously unrecognized pattern in plasmids across prokaryotes, that could lead to new insights into the evolutionary role that plasmids play.

## Introduction

Plasmids are an important source of prokaryotic genetic variation, though they play different roles for different clades. In the pathogenic members of the *Enterobacteriaceae*, for instance, plasmids are highly variable and provide their hosts with the opportunity to acquire new genes. In *Borrelia*, on the other hand, the number of plasmids and their genetic content seem far more consistent between strains (1). Still other prokaryotes rarely carry plasmids. These different plasmid roles have important effects on the evolution of these organisms, including the separate pathways by which they can become resistant to antibiotics. Many recently emerged antibiotic resistance genes are primarily found on plasmids, including cephalosporinases such as CTX-M, carbapenemases such as KPC, aminoglycoside acetylases such as aac6’-1b (2), and the colistin resistance gene MCR-1 (3). Examples of these genes with perfect sequence identity have been found in different plasmid backbones and incompatibility groups. Though many individual plasmids have been studied in detail, large scale studies of plasmid populations are required to understand the effect of different plasmid roles on prokaryotic evolution and behavior.

In several studies of sets of plasmids, researchers have noted plasmids that appear to be ‘mosaics;’ in other words, they contain genes from several ancestral genetic sources, likely due to recombination (though the exact definition of mosaic depends on the analysis) (1, 4–7). Because the effects of recombination are highly variable, even in controlled settings (8), the study of mosaic plasmids has been largely observational, with scope restricted by the number of plasmid sequences available. In 2011, Bosi et al. (9) identified putative ‘atypical’ genes in the set of plasmid sequences available at the time, using the parametric methods of GC content and dinucleotide profiling. They found such genes to be widespread, and in certain cases could determine that this ‘atypical’ signature was due to transfer of genetic material to the plasmid from another source, demonstrating that these were mosaic plasmids. For 75.7% of the genes for which the source could be found, the source molecule was another plasmid, suggesting frequent genetic exchange between plasmids.

Many of the known examples of mosaic plasmids are associated with the actions of transposases, particularly when antibiotic resistance genes are on the mosaic fragment (7, 10). Transposons are found in high abundance on plasmids, and some transposases, such as the IS26 transposase, have been observed almost exclusively on plasmids due to their mechanism of transposition (4). Because IS26 causes large scale rearrangements, it decreases the stability of the genetic element it is found on, and could delete essential genes when present in chromosomes. Additionally, target sequence can affect transposon localization, such as for IS26, which seems to target some other transposases for insertion (11). Other recombination processes have been shown to contribute to mosaic plasmid formation, including homologous recombination (11) and non-homologous end joining (1). Homologous recombination between sites on a single plasmid has been detected in cultures grown from single isolates (12), suggesting that mosaic plasmids form frequently when a recombination method is available. Mosaic plasmids have often been described that expand genes that enhance the fitness of their hosts (7), but recombination that stabilized an invasive plasmid has also been observed in a model experiment (13).

Each individual mosaic plasmid can accumulate many fitness-enhancing genes and provide great benefit to its host organism, as seen with the emergence of large, multi-drug resistance plasmids in the *Enterobacteriaceae* (14), but that mosaic plasmid can also be costly to maintain. Many factors can potentially influence the formation of mosaic plasmids, such as transposon abundance, variability of selection pressures, incompatibility groups, abundance of conjugative plasmids, and host tolerance for foreign genetic material. These factors vary across prokaryotic taxa and plasmid populations. Each of these factors has tradeoffs for the host organism, adding to the overall tradeoff of having a highly adaptable genome versus a stable genome. The fraction of mosaic plasmids in the plasmid population is likely to evolve within populations over time due to both variation and potential selective pressures. Thus, it might be expected that the fraction of mosaic plasmids from a prokaryote’s plasmid population is reflective of its evolutionary history and predictive of its response to new pressures.

Previous studies identifying mosaic plasmids have used gene synteny (1), parametric measures (9), or sequence identity (6, 7) comparisons. Gene synteny can be a powerful tool in instances where there have been relatively few rearrangements, but could lead to false positives in systems with frequent rearrangements and high overall genetic diversity. Synteny can also require functional knowledge of the genes involved to determine the level of identity necessary to declare genes homologous. For these reasons, synteny is not well suited to large studies across highly diverse sets of plasmids.

Parametric measures of gene composition can be applied on a large scale, and are less affected by database biases. They are also prone to error, since GC content and dinucleotide profiles vary even for genes that have been vertically transferred for many generations. This is particularly true for comparisons to smaller genetic elements (since fewer genes are available to establish the proper baseline), increasing the false positive rate. Parametric methods also suffer from false negatives, missing mosaic events where plasmids from similar genetic backgrounds have exchanged DNA.

Sequence identity can be specific enough to confidently predict that a DNA sequence has transferred between genetic contexts (15), but can only be used to identify transfer events that have happened recently (i.e. with few or no subsequent point mutations). Sequence identity comparisons thus have low rates of false positives but high rates of false negatives, and are most effective in identifying populations where DNA transfer is frequent relative to the base mutation rate.

In the research described below we use a sequence-identity-based, quantitative definition of ‘mosaic’ to study mosaic plasmids across 8,592 plasmid sequences from the RefSeq plasmid database maintained by the National Center for Biotechnology Information (hereafter referred to as the NCBI plasmid database). Using our quantitative, universal definition of mosaic plasmids, we compare the fraction of plasmids that, using these conservative criteria, are mosaic across different plasmid populations. By extending the concept of mosaic plasmids to the population level, we can begin to ask questions about the forces that favor or disfavor mosaic plasmids, and the implications of mosaic plasmids for important prokaryotic traits such as the spread of antibiotic resistance. Our initial hypothesis is that mosaic plasmids would be found across many prokaryotic clades based on the fundamental mechanisms involved, and because they have been previously observed in species from the *Enterobacteriaceae* (5, 7), *Rhizobium* (12), and *Borrelia* (1) clades. We also hypothesize that mosaic plasmids would be most abundant in the *Enterobactericeae*, where mosaic plasmids have been most frequently observed and contribute to high levels of antibiotic resistance(6).

## Results

### Abundance of mosaic plasmids

For our analyses, we used all 8,592 finished or draft prokaryotic plasmid sequences in the NCBI plasmid database (Table 1) except where noted otherwise. We define a plasmid as mosaic if it contains a region of 500 bp or longer with 100% identity to the sequence of another plasmid (called a mosaic fragment), and where those two plasmids have less than 93.90% global sequence identity (see *Methods* section for details). For the purposes of this study, we will refer to plasmids that pass our criteria as “mosaic” and those that do not pass as “non-mosaic”.

**Table 1:**
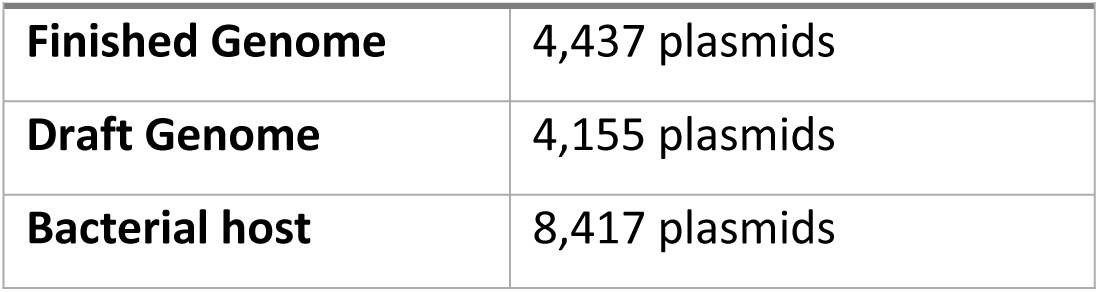
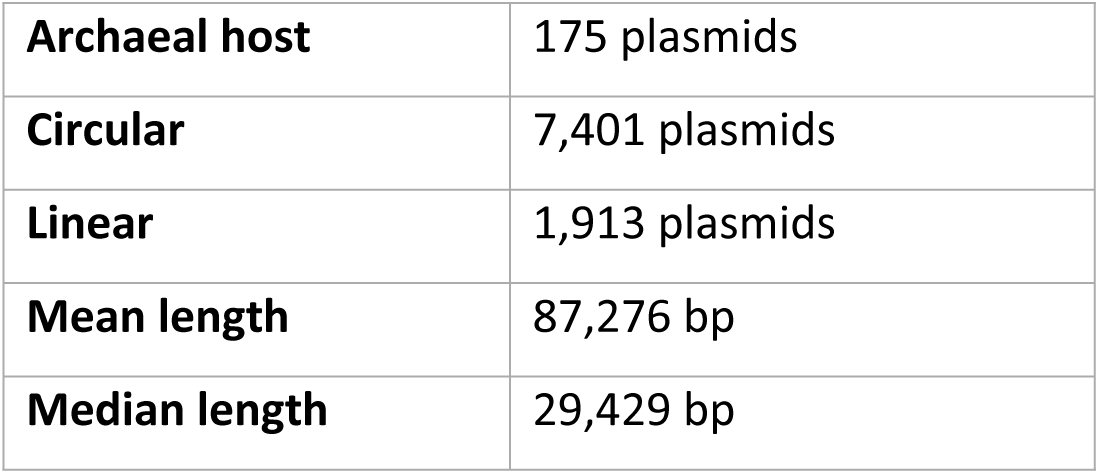
NCBI plasmid database properties.

| Table 1: NCBI plasmid database properties. |  |
| --- | --- |
| Finished Genome | 4,437 plasmids |
| Draft Genome | 4,155 plasmids |
| Bacterial host | 8,417 plasmids |
| **Archaeal host** | 175 plasmids |
| **Circular** | 7,401 plasmids |
| **Linear** | 1,913 plasmids |
| **Mean length** | 87,276 bp |
| **Median length** | 29,429 bp |

Applying our definition to the NCBI plasmid database, we found that 46.35% of plasmids are mosaic. We used network analysis to identify particular traits of mosaic plasmids and the DNA shared between them, where we term each instance of shared DNA as a mosaic fragment, represented as a line connecting mosaic fragments (Figure 1A, only mosaic fragments over 5kb are shown). Most mosaic plasmids share identity with other plasmids over less than 30% of their full length (median = 26.93%), though over 600 mosaic plasmids share identity over more than 95% of their length (Figure 1B). Of the links between mosaic plasmids, 82.71% are between plasmids isolated from different species (Figure 1C). Each mosaic plasmid can have between 1 and several thousand links (median = 12, Figure 1D). A single plasmid, pMR0716_tem1 (NZ_CP018104) isolated from *Escherichia coli*, was identified with 20,423 links. Though some mosaic plasmids have links to hundreds of other plasmids, the vast majority are connected to fewer than 25 (median = 8, Figure 1E). Only 1.63% of mosaic fragments are greater than 5,000 bp, but these longer mosaic fragments are still found in 19.16% of plasmids from the NCBI plasmid database. Of these longer mosaic fragments, 76.97% are shared between plasmids isolated from different species.

**Figure 1:**
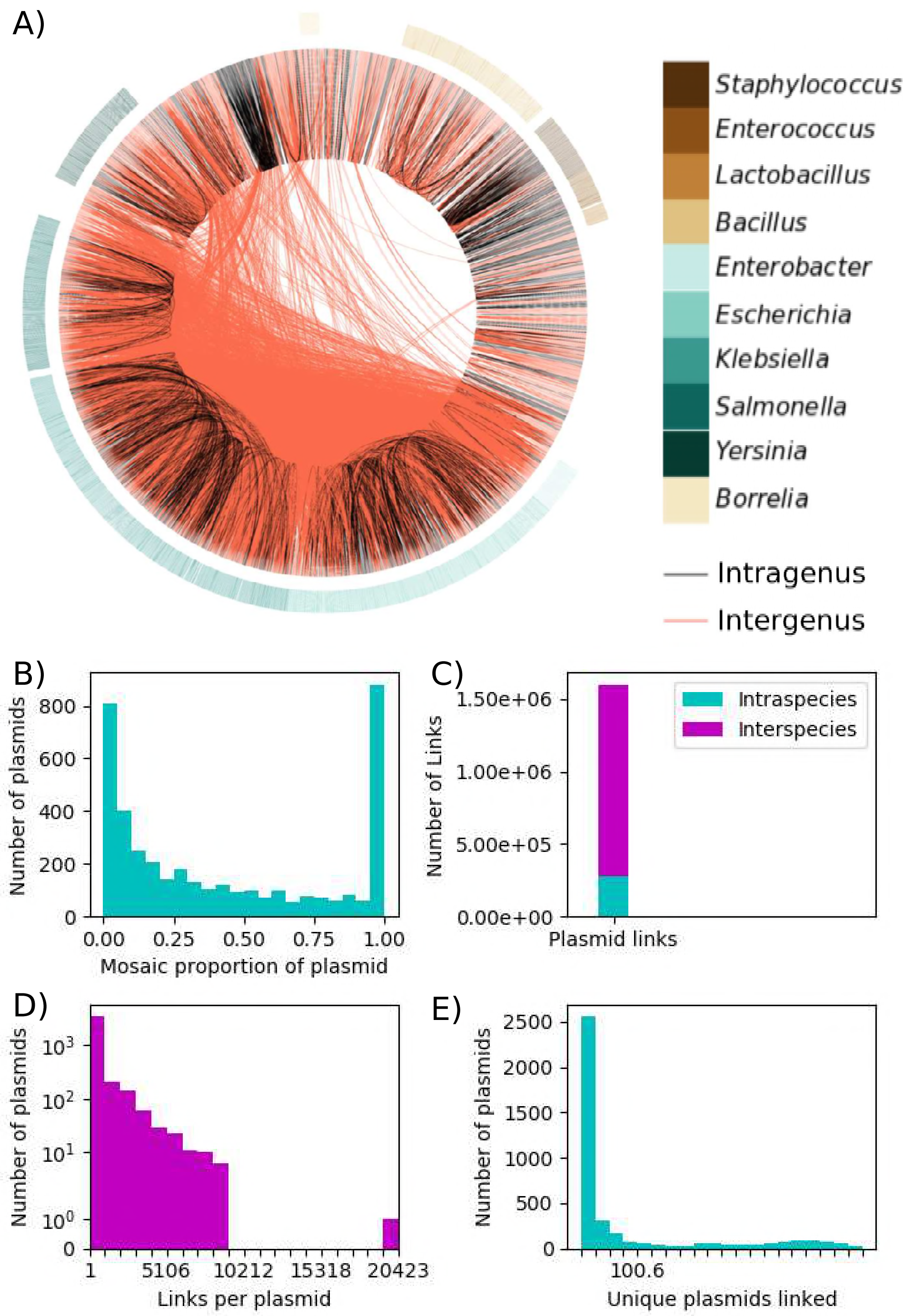
Properties of mosaic plasmids. (A) Network diagram of mosaic fragments shared by mosaic plasmids. Lines connecting two points on the circle represent a shared fragment of 5 kilobases or more. A black line indicates that the connected plasmids were isolated from the same species, and red lines indicate that they came from different species. Plasmids are ordered by the taxonomy of the prokaryote from which they were isolated. Colored lines extending away from the circle highlight selected genera. We generated the figure with Circos (28). (B) Histogram of the distribution of mosaic plasmids by the proportion of the plasmid covered by mosaic fragments. (C) Histogram of the distribution of mosaic plasmids by the number of other plasmids they are linked to. Full range of links per plasmid was divided into 20 even bins. Y-axis scale is symmetric log. (D) Stacked bar chart of mosaic plasmids with links to plasmids isolated from the same species or isolated from a different species. (E) Histogram of unique other plasmids that each mosaic plasmid shares a mosaic fragment with.

Mosaic plasmids in the database tend to be longer than non-mosaic plasmids, with an average length of 120,682 bp and median length of 62,653 bp, compared with an average length of 61,926 bp and median length of 13,415 bp for non-mosaic plasmids (Figure S1). These numbers are likely influenced by our strict requirements for mosaic plasmids, and by the fact that most known megaplasmids (greater than 1 Mbp) are mosaic.

### Distribution of mosaic plasmids over incompatibility groups

We were able to identify incompatibility groups for only 1,890 plasmids using the PlasmidFinder database (16). In these plasmids, we found an average of 1.39 rep genes. Where multiple rep genes were present, we classified the plasmid according to the highest confidence match to PlasmidFinder. Most of these plasmids fit our definition of mosaic (Figure 2A) and many had exchanged mosaic fragments with plasmids from other incompatibility groups (Figure 2B).

**Figure 2:**
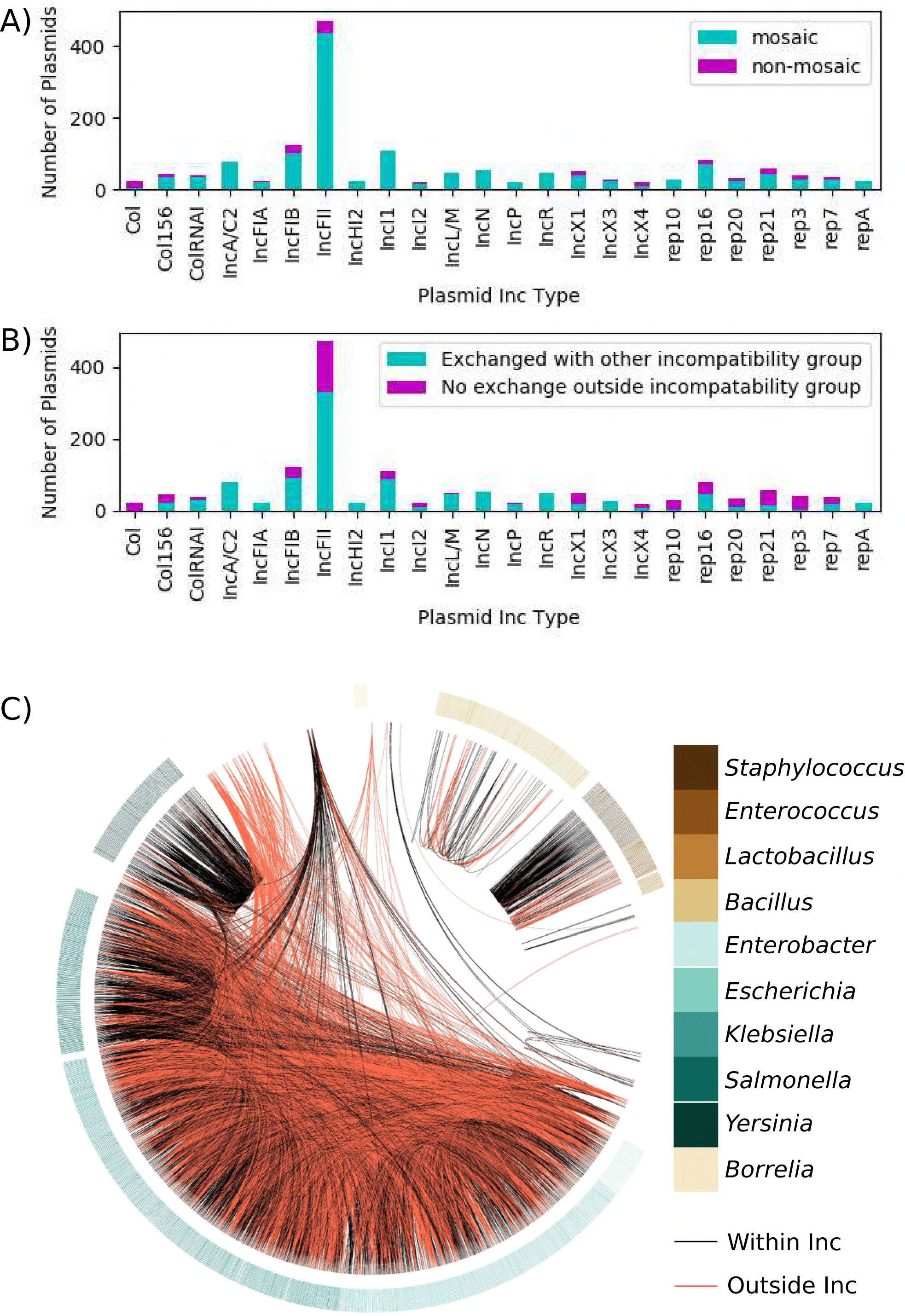
Mosaic plasmids by Inc group. (A) Stacked bar chart of mosaic and non-mosaic plasmids detected within the top 20 incompatibility groups by frequency in this dataset. (B) Stacked bar chart of mosaic fragment sharing within and between incompatibility groups for the same incompatibility groups as in (A). (C) Network diagram of mosaic fragments shared where incompatibility group could be identified for both plasmids. A black line indicates that the connected plasmids are from the same incompatibility group, and red lines indicate that they came from different incompatibility groups. Plasmids are ordered by the taxonomy of the prokaryote from which they were isolated. Colored lines extending away from the circle highlight selected genera. We generated the figure with Circos (28).

We again created a network to visualize the sharing of large (> 5 kbp) mosaic fragments between plasmids from the same or different incompatibility groups (Figure 2C). This visualization shows that some of the mosaic fragments shared over large phylogenetic distances stay within the same incompatibility group, as well as a bias in *Yersinia* for exchange within an incompatibility group. 71 of the 158 Yersinia plasmids are IncFII, including several variants of the *Y. pestis* plasmid pCD1.

### Distribution of mosaic plasmids over genera

Since our definition of mosaic is based on comparisons within the database, we hypothesized that as the number of sampled plasmids in a population increases, the proportion of those plasmids identified as mosaic would also increase. This hypothesis implicitly assumes that the underlying distribution of mosaic plasmids is uniform across all plasmid populations. We tested this hypothesis on all genera with at least ten sequences in the NCBI database, equaling 8,272 plasmids across 117 genera. Our analysis does not support our hypothesis, as there is no apparent correlation between the number of plasmid sequences in the database from an organism and the fraction of those plasmids deemed to be mosaic (Figure 3, Table S1). The three *Enterobacteriaceae* genera with the highest number of plasmid sequences had similar percentages of mosaic plasmids in our analysis - *Escherichia* (901 plasmids, 66% mosaic), *Klebsiella* (598, 73%), and *Salmonella* (364, 66%). The gram-positive *Staphylococcus* (476 plasmids, 57% mosaic) and *Bacillus* (533, 55%) also have majority mosaic plasmid populations. In contrast, the plasmid populations isolated from *Lactobacillus* (357 plasmids, 30% mosaic) and *Borrelia* (995, 5% mosaic) have low percentages of mosaic plasmids. The unique characteristics of the *Borrelia* plasmids, including high plasmid carriage, stable plasmid repertoire, and high abundance of linear plasmids, may complicate comparisons with other plasmid populations. Plasmid populations of all other genera include fewer than 200 plasmids and range from 0% to 100% mosaic. The only genus with 100% mosaic plasmids is *Piscirickettsia*, with 61 sequenced plasmids.

**Figure 3:**
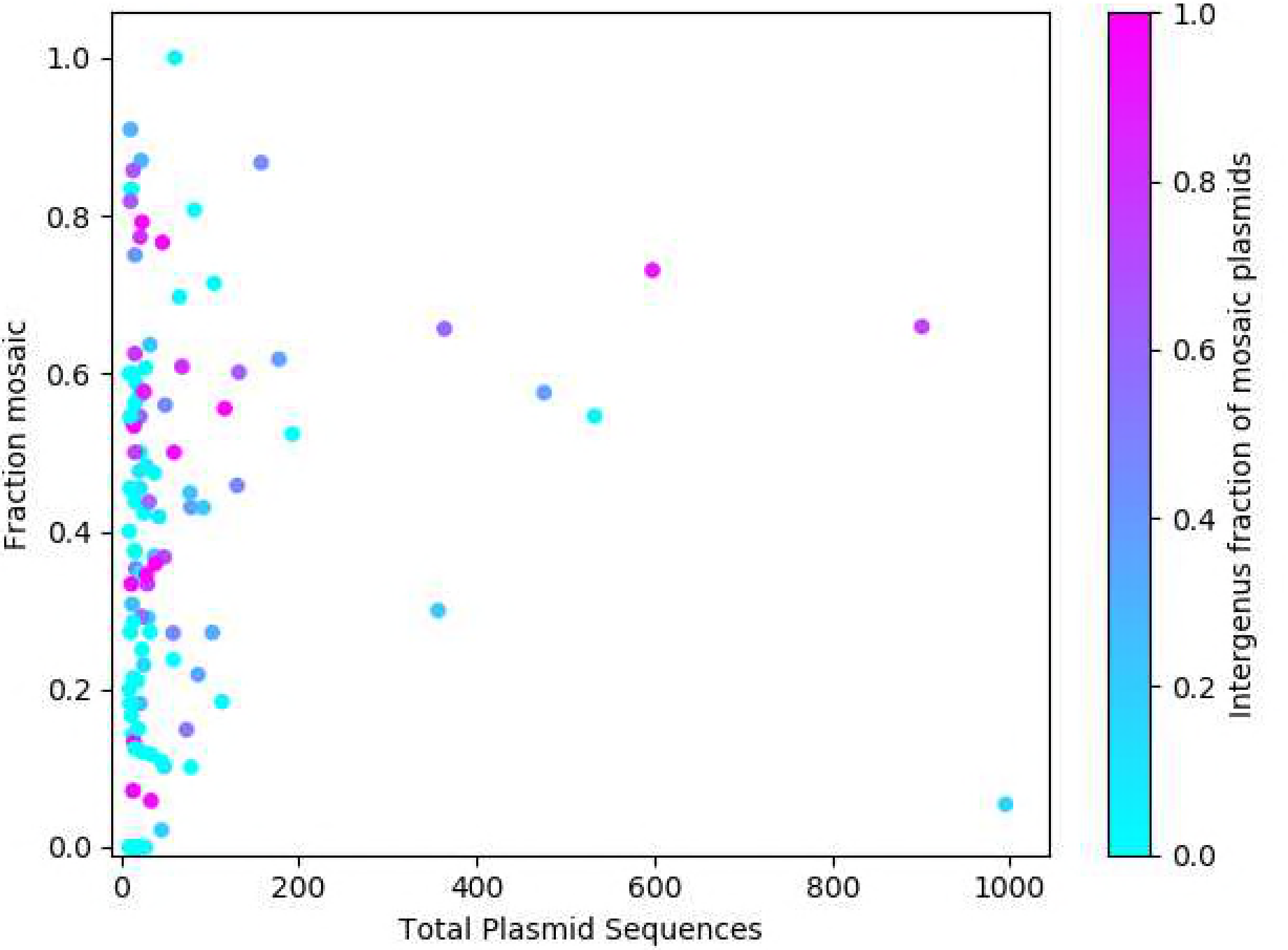
Scatter plot of total plasmid sequences isolated from a particular genus by the fraction of those plasmids that are mosaic. Points are colored by the fraction of mosaic plasmids from that genus that share at least one mosaic fragment with a plasmid isolated from another genus.

Many of the detected mosaic plasmids shared mosaic fragments with plasmids isolated from other genera (Figure 3). This was especially clear in the *Enterobacteriaceae*, potentially due to the high sampling in that family. Plasmid populations isolated from *Piscirickettsia* and *Bacillus*, despite their high fractions of mosaic plasmids, had few connections to plasmids from other genera in the database: 3.28% and 4.12% of their respective mosaic plasmids contained such a connection. Similarly, of the 101 mosaic plasmids isolated from *Rhizobium* species (52.33% of the 193 total *Rhizobium* plasmids), none shared a mosaic fragment with plasmids from any other genus.

### Mosaic plasmid characteristics

There are many potential factors that could affect the prevalence of mosaic plasmids within different plasmid populations. We explored two main hypotheses: 1) transposases increase the rate of mosaic plasmid formation, and thus mosaic plasmids will be enriched for transposases compared to non-mosaic plasmids; and 2) antibiotics create selection pressure for antibiotic resistance gene spread via mosaic plasmids, and thus mosaic plasmids will be enriched for antibiotic resistance genes compared to non-mosaic plasmids.

To test our transposase hypothesis, we identified transposases using a Hidden Markov Model (HMM) transposase database developed from the J Craig Venter Institute Genome Property, GenProp1044 (17). After identifying transposases, we calculated the proportion of genes on each plasmid that were transposases (to eliminate bias resulting from mosaic plasmids greater median length). We then compared the distribution of transposase gene proportions in mosaic and non-mosaic plasmids. For this analysis we only considered plasmids from finished genomes with annotated coding regions (n = 4,055). We found that mosaic plasmids are composed of a significantly higher proportion of transposases (median = 5.0% of genes) compared to non-mosaic plasmids (median = 0.0% of genes), with p = 6 × 10^−195^ (two-sided Mann-Whitney U test), supporting our hypothesis (Figure 4A). Despite the high abundance of transposases in mosaic plasmids, we could not detect any transposases in 389 of the finished plasmids (n = 1560) we defined as mosaic. Also, 765 non-mosaic plasmids do have a transposase.

**Figure 4:**
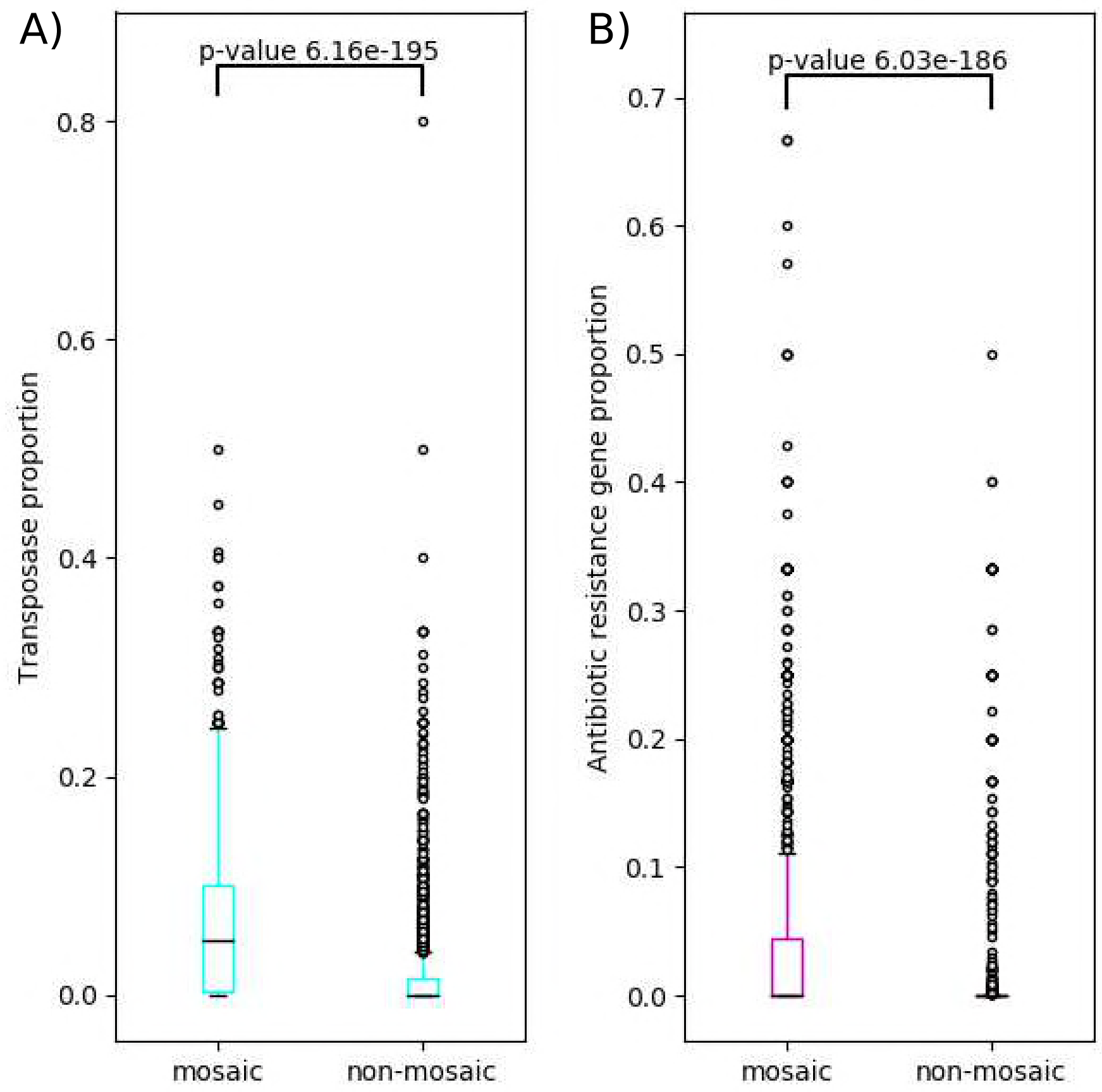
Boxplot of mosaic and non-mosaic plasmids over the proportion of their genes of interest. Bottom and top of the box represent the first and third quartiles, respectively. Black line represents distribution median. Whiskers extend out 1.5 times the interquartile range. Black circles represent individual outlier data points. P-values calculated using Mann-Whitney U two-tailed test with the null hypothesis of equal distributions. (A) Transposases and (B) Antibiotic resistance genes.

As a complimentary method, we identified inverted repeats (IR) using EMBOSS(18) as signs of potential transposon or insertion sequences in our mosaic plasmids. Table 2 shows where IR occurred within 10 bp of the boundaries of mosaic fragments in each of their host plasmids. The 10 bp gap was included because we expect the mosaic boundary determination to vary in some cases due to the random nature of the surrounding plasmid DNA. We found that few mosaic appear to be simple transposons - flanked by IR and containing a transposase - with 64.12% of mosaic fragments not flanked by IR, and 43.42% not containing any transposase genes.

**Table 2:** Mosaic fragment correlation with inverted repeats

| **Table 2: Mosaic fragment correlation with inverted repeats** |  |  |
| --- | --- | --- |
| **Condition** | Percent of mosaic fragments | Percent also containing $\geq 1$ transposase |
| **All mosaic fragments** | 100.00% | 56.58% |
| **Flanked by matching IR** | 0.91% | 0.65% |
| **Flanked by unmatched IR** | 1.96% | 0.84% |
| **Flanked by only one IR** | 33.02% | 17.97% |
| **Not flanked by any IR** | 64.12% | 36.47% |

We identified resistance genes using the Resfams HMM database (19), and calculated the proportion of genes on each plasmid that confer antibiotic resistance, again limiting to plasmid sequences containing at least ten genes (Figure 4B). We found that mosaic plasmids (median = 0.0% of genes) contain significantly more antibiotic resistance genes non-mosaic plasmids (median = 0.0% of genes), with p = 6 × 10^−186^ (two-sided Mann-Whitney U test), supporting our hypothesis.

### Analysis of highly shared mosaic fragments

We next turned to the question of the identity of mosaic fragments shared between plasmids, starting by identifying the number of occurrences of the mosaic fragments in the full NCBI plasmid database. For this analysis, mosaic fragments must be greater than 500 bp long and present in more than one plasmid. To focus on the mosaic fragments that are shared most frequently, in cases where one mosaic fragment was a subsequence of another, the larger mosaic fragment was reduced to the subsequence and combined with the smaller mosaic fragment. The identified fragments range in size from 501 bp to 55,524 bp, with a median length of 1,146 bp. We enumerated the number of occurrences of each mosaic fragment in the NCBI plasmid database, counting instances where a mosaic fragment is present in multiple locations on the same plasmid (Figure 5A). We found that the clear majority of mosaic fragments occur in fewer than 250 locations in the NCBI plasmid database, but that there are a few highly abundant outliers (Table S2). Only 35.22% of mosaic fragments contain any part of functional transposase sequence, and these fragments are similarly distributed to the mosaic fragments without transposases (Figure 5B). It is possible this is an artifact of our requirement that sequences be 100% identical to be considered the same mosaic fragment; for instance, if a mosaic fragment is moved as a transposon and the transposase undergoes one or more point mutations while adjacent sequences do not, it would not be considered part of the same mosaic fragment. If we relax the mosaic definition to allow 99%-identical sequences to be considered mosaic fragments, then the percent including some transposase sequence increases to 35.69% (Figure S2).

**Figure 5:**
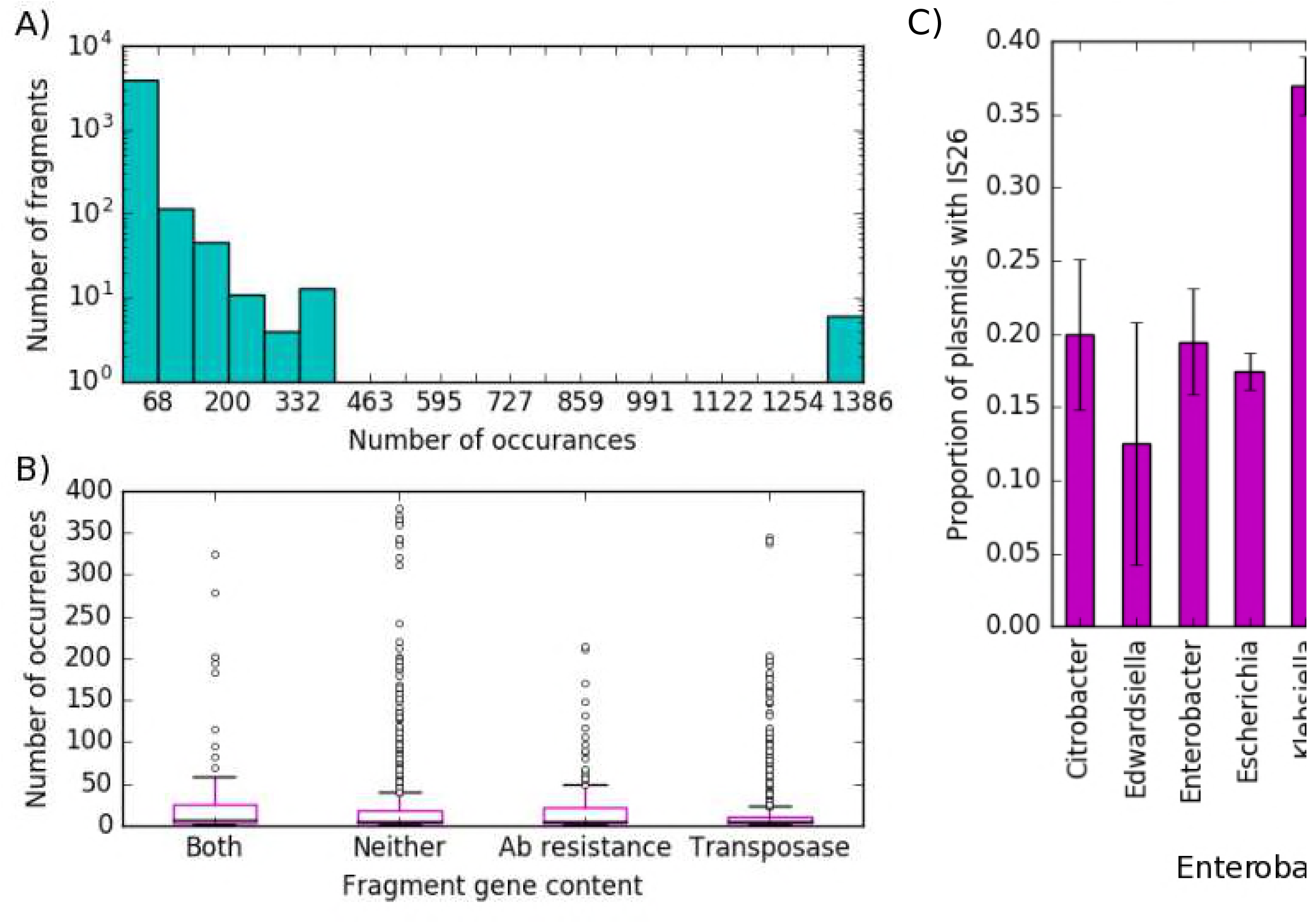
Mosaic fragment abundance, composition, and distribution. (A) Histogram of DNA fragments of at least 500 base pairs over their number of appearances (at 100% nucleotide identity) in the NCBI plasmid database. Occurrences are grouped into 20 even bins across the full range of data. Y-axis is log scale. (B) Boxplot comparing the distributions of mosaic fragments by gene content. Boxes composed as in Figure 3. The six outlier points have been removed to make the distributions visible. (C) Histogram of the proportion of plasmids in each genus that contain the IS26 transposase gene. Only genera with at least one plasmid containing the IS26 transposase gene are included. Error bars represent standard error of the proportion. X-axis is sorted by host organism family, though only the *Enterobacteriaceae* are represented by more than one genus.

Examining specific sequences revealed that the six mosaic fragments with between 1,300 and 1,400 occurrences all contain part of the IS26 transposase (in one fragment the transposase is annotated as incomplete). These mosaic fragments often co-occur, but each occurs separately in at least one location in the NCBI plasmid database (information on genetic context for these fragments in Table S2). This means that the IS26 transposase is found in the NCBI plasmid database more frequently than the most abundant mosaic fragment (1,386). To determine if the IS26 transposase is widely distributed, or highly abundant in certain taxa, we assessed the proportion of plasmids containing the IS26 transposase from each genus represented in the NCBI plasmid database (Figure 5C). This analysis demonstrates that IS26 is phylogenetically constrained, since each genus that contains the IS26 transposase in greater than 5% of its plasmids, apart from *Photobacterium* (which had 7 plasmids present in the NCBI database), is a member of the *Enterobacteriaceae*. A BLAST of the IS26 protein sequence against pMR0716_tem1, the most highly linked plasmid sequence, revealed 37 complete copies (and at least one incomplete copy) of the transposase.

To determine if there were any mosaic fragments present in more diverse contexts than the IS26 transposase, we identified the number of distinct genera carrying each mosaic fragment (Figure S3). We found that there are mosaic fragments found in plasmids from up to 24 genera, compared to 19 genera for the IS26 transposase. Each of the diverse context mosaic fragments was still found in *Entrobacteriaceae* plasmids, particularly from the most well-sampled genera - *Escherichia, Klebsiella*, and *Salmonella*.

## Discussion

### Mosaic plasmids are highly abundant and unevenly distributed across plasmid populations

Based on our results, mosaic plasmids appear to be very common across different plasmid populations. Our conservative definition of mosaic allowed us to be confident in our positive results across diverse plasmid types, but it also missed large groups of plasmids that may be considered mosaic in a more general sense. First, by basing our definition on comparison to other known plasmid sequences, from a database biased towards human pathogens and their antibiotic resistance determinants, we missed mosaic plasmids that share mosaic fragments with yet-unsequenced plasmids. Second, we have excluded the contribution of other mobile DNA elements, such as integrative conjugative elements and phage, to focus on DNA movement within plasmid populations. Third, our analysis missed any mosaic plasmids that shared a mosaic region with a chromosome rather than a plasmid. Fourth, it excludes potential mosaic fragments that have undergone subsequent point mutations after they were shared between plasmids, though these have been called mosaic plasmids previously (1).

The frequency of each of these events (except for the first) in a plasmid population represents a property of that population, and likely plays an important role in shaping plasmid and prokaryotic evolution. Bosi *et al*. demonstrated that exchange between plasmids is most common on a global level (9), but types of exchange may vary greatly from population to population. Despite excluding other types of exchange, we found that over 40% of plasmid sequences from the NCBI plasmid database fit our definition of mosaic, suggesting that plasmid sequence and identity are very fluid and dynamic in nature.

We attempted to account for the first case listed above - that true mosaic plasmids were missed because of low sampling in some plasmid populations - by comparing the mosaic fraction for plasmids isolated from each prokaryotic genus against the number of plasmids isolated from that genus. The lack of a strong correlation suggests that sampling depth is not the primary driver of mosaic fraction, and that, despite their overall prevalence, mosaic plasmids are unevenly distributed across prokaryotic taxa. The two hypotheses we tested - that transposases and antibiotic resistance genes are enriched in mosaic plasmids - were each supported. However, as seen in Table 2, and Figures 4 and 5B, they are incomplete explanations for the formation of mosaic plasmids. Further, they are not independent variables. Figure 5B demonstrates that mosaic fragments exist with antibiotic resistance genes and not transposases and vice versa, but their frequent co-localization likely increases the frequency of mosaic events around both types of genes.

We suggest that transposases represent a mechanism of variation and that antibiotic resistance genes represent a selective pressure for the formation of mosaic plasmids. A third potential component is exposure to novel genetic material, with conjugation representing one mechanism. We did not explore this possibility because we were concerned that we could not accurately identify this trait across the diverse plasmids from this dataset, but conjugation is a prime target for future, more focused studies.

Our results indicate that alternative mechanisms and selection pressures are necessary to account for the full range of mosaic fragments. Accounting for these alternatives would improve our understanding of plasmid populations dynamics. Moreover, the study of plasmid populations can be used to create hypotheses about genetic and evolutionary forces. For instance, identifying common mosaic fragments in plasmids isolated from a prokaryotic clade could be used to identify likely fitness-enhancing genes within that clade.

Non-genetic factors also probably play a role in the distribution of mosaic plasmids. HGT of some sort is required for mosaic plasmid formation, and previous studies have shown that shared environment is the most important factor in promoting HGT(15). Like any type of genetic variation, mosaic plasmids will be more favored in some environments than others. Similarly, ecological niches could influence the favorability of mosaic plasmids. Interestingly, plasmids from *Yersinia pestis* (71% of plasmids from the *Yersinia* genus) and *Borrelia* are both human pathogens whose lifecycles include insect vectors. Yet 98% of *Y. pestis* plasmids are mosaic, while only 5% of *Borrelia* plasmids are. While genetic factors likely play a role in this difference, it would be interesting to determine if specific environmental factors also influence mosaic fraction. Studies on systems such as this would improve our basic understanding of prokaryotic variation. Understanding ecological interaction with mosaic plasmids would prove invaluable in the study of environments such as hospital surfaces, where human action shapes the community structure and plasmid exchange can have direct health impacts. We were unable to incorporate plasmid host source environment or lifecycle in our study design, in part because the source environmental information is not consistently coded in the NCBI plasmid database. The rewards of understanding how environment and ecology affect mosaic plasmid creation warrant studies focused on those questions.

### Mechanism of mosaic plasmid formation

Our analysis suggests that mosaic plasmids are a broader phenomenon than transposon hopping between plasmids through three lines of evidence. First, other mechanisms for mosaic plasmid formation have been observed previously, including homologous recombination (11), non-homologous end joining (1), and large scale rearrangements of unknown mechanism (7). Of these, homologous recombination has the potential to play a major role in plasmid populations where some mosaic fragments are already highly abundant. Second, Figure S1 shows over one hundred highly abundant mosaic fragments that have no overlapping transposase sequence, even when only 99% identity is required to call them mosaic. These fragments may be adjacent to transposases in some but not all plasmids, suggesting that either distinct transposases are moving the same mosaic fragments or other mechanisms are involved. Third, several of the mosaic fragments are larger than would be expected from transposition events alone. Further studies will be required to definitively determine the contribution of each of these to mosaic plasmid formation and whether those contributions vary along with mosaic fraction across plasmid populations. Regardless of the extent to which transposases are responsible for mosaic plasmids, it is interesting to understand why the distribution of mosaic plasmids is highly variable across taxa, and to know which transposases (beyond the IS26 transposase) support intergenus transfer events.

### Organisms with noteworthy plasmid populations

Three plasmid populations stood out in our analysis for different reasons: plasmids from *Borrelia*, plasmids from the *Enterobacteriaceae*, and plasmids from *Piscirickettsia*. The plasmid population of the *Borrelia* genus, members of which can cause Lyme disease or relapsing fever, was a prominent outlier in Figure 2, having both many plasmid sequences (995) and a low mosaic fraction (5%). Previous analysis has determined that these plasmids do undergo recombination, leading to mosaic plasmids (1); however, these events were determined to happen at a lower rate than synonymous point mutations. In fact, this study used mosaic plasmids as stable genetic markers. The difference in methodology between our analysis and this previous analysis thus reveals that the *Borrelia* plasmid population forms mosaic plasmids much less frequently, relative to point mutation rate, than most other plasmid populations. Given the high density of plasmids in *Borrelia* species, it is surprising that they do not form mosaic plasmids more frequently. We found that the *Borrelia* plasmid population contains few transposases or antibiotic resistance genes, a finding that could offer some explanation for their stability. In comparison to the mosaic plasmids identified by our definition, *Borrelia* plasmids may be under very different evolutionary pressures.

In contrast to *Borrelia*, much of what made the genera of the *Enterobacteriaceae* family interesting could be attributed to their abundance in the NCBI plasmid database, rather than the properties of their plasmid population. Many of the previous examples of mosaic plasmids in the literature come from *Enterobacteriaceae* (5–7), and the pathogenic strains are frequently multidrug resistant (14, 20). Despite this coincidental support for a high mosaic fraction, the *Enterobacteriaceae* were surpassed in mosaic fraction by several genera, including *Yersinia* with 87% mosaic plasmid sequences (out of 158 total), and *Piscirickettsia* with 100% mosaic plasmid sequences (out of 61 total). Though *Enterobacteriaceae* plasmids are not outliers relative to other plasmid populations, their mosaic plasmids are still important. For instance, the prominence of the IS26 transposase in Figure 3A is due to the high representation of *Enterobacteriaceae* plasmids in the database. Its frequency in those plasmids means that it is a potential target for diagnostics or treatments designed to limit the spread of resistance genes in pathogens from the *Enterobacteriaceae* family. It is unclear from this analysis whether other transposases are as prominent in other families. To determine if attributes of *Enterobacteriaceae* plasmids are unique, it will be necessary to obtain a substantial number of diverse plasmid sequences from other prokaryotic families to make a comparison.

The plasmids from *Piscirickettsia* are interesting because they are the only genus with 100% of its plasmids identified as mosaic in our analysis. *Piscirickettsia*, known as salmon pathogens, can be exposed to high levels of antibiotics when present in salmon farms. Recent work has determined that antibiotic resistance genes found in fish stock pathogens can be traced to fish feed (21). The *Piscirickettsia* plasmids present in the NCBI plasmid database come from strains isolated from salmon farms in southern Chile across multiple studies by two research groups over several years (22, 23). Since these plasmids came from one geographic region, they may not represent the global diversity of *Piscirickettsia* plasmids - but even if these plasmids had all come from a single study the mosaic fraction would be notably high. Interestingly, among plasmid populations with a high mosaic fraction, the *Pisciricksettsia* plasmids were unusual in that all but two of their plasmids shared mosaic fragments only with other *Piscirickettsia* plasmids. There may still be plasmids from other organisms that introduce these mosaic fragments, which then spread throughout the *Piscirickettsia* plasmids, but if so then these source plasmids have not yet been sequenced.

Interestingly, the 206 archaeal plasmids included in the plasmid database did not stand out in this analysis of mosaic character. 37.5% of plasmids from the most common archaeal genus of origin, *Haloferax*, were mosaic, with no intergenus links - an identical profile to the bacterial genus *Geminocystis* (Table S1). While further sampling and a more direct focus may reveal unique features of archaeal plasmid population dynamics, for now they appear to fall within the range of bacterial plasmid populations.

### Challenges for future study of mosaic plasmids

This study highlights the power of utilizing a resource like the NCBI plasmid database to answer population level questions; however, its biases and composition make it inappropriate to address certain questions. Many of the new questions raised by our analysis will require new research studies specifically designed to address target hypotheses. Confidently separating plasmid and chromosomal sequence from short-read sequencing data is challenging (24), and both physical separation and long-read sequencing are expensive and time consuming. Mosaic plasmids are particularly challenging: their high carriage of repetitive DNA elements, including transposons, frustrates short-read *de novo* assembly, and their mosaic nature confuses reference-based software. Model systems can be used to study plasmid dynamics for specific populations, but study design incorporating highly variable events such as mosaic plasmid formation will be challenging. Given the quantity of further questions in plasmid population dynamics, further database mining, targeted sequencing projects, and model systems will each be required depending on the specific questions.

## Methods

### Plasmid sequences

All scripts and instructions to reproduce these analyses and figures are provided in the GitHub repositories referenced below. We acquired all plasmid sequences from the NCBI Refseq database on April 10^th^ 2017 (ftp://ftp.ncbi.nlm.nih.gov/refseq/release/plasmid/). We then concatenated the plasmid nucleotide sequences from the Refseq release into a single FASTA file (ncbi_plasmid.fna), and removed all sequences of plasmids isolated from Eukaryotes (38 sequences removed). Known incomplete sequences (accession numbers starting with prefixes other than “NC” or “NZ”) were removed with custom scripts. We converted the remaining sequences to a BLAST database using BLAST command line tools version 2.2.31+ and the command ‘makeblastdb -in ncbi_plasmid.fna -dbtype nucl -out ncbi_plasmid’. We extracted plasmid protein sequences from the Genbank file using a custom script. We derived a BLAST database from this the command ‘makeblastdb -in ncbi_plasmid.faa -dbtype prot -out ncbi_plasmid_proteins’.

### Mosaic plasmid definition

The definition used for mosaic was plasmids that have a region of at least 500bp with 100% nucleotide identity with another plasmid, where those two plasmids have less than 93.90% identity over their full lengths. We chose 500 bp based on a previous definition of horizontal gene transfer (15) and because a minimum was required that could not easily arise by chance. At 93.90% identity, assuming a random distribution of point mutations, the probability that two 500,000bp plasmids have 500bp at 100% identity (probability of false positive, P(fp)) is <10^−8^. Since there are fewer than 10^4^ plasmids in the NCBI database, there are fewer than 10^8^ possible pairwise comparisons between them. Thus, even if all of the plasmids in the database were highly similar, we would expect fewer than one false positive mosaic plasmid identification using this threshold. See Eq. 1 below where N = estimated plasmid length in bp, M = number of identical bases between the two plasmids, and X = length of 100% identity region.

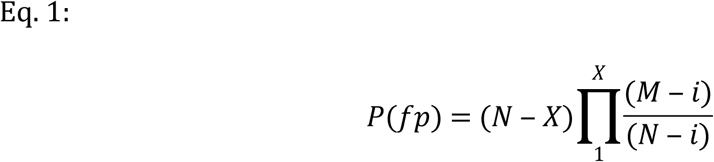

We identified plasmids putatively meeting this definition using BLAST with the command ‘blastn -query ncbi_plasmid.fna -db ncbi_plasmid -perc_identity 100 - outfmt 6 -num_threads 8 -out perfect_plasmid_matches.txt’. We then used a Python script (Python 3.5.2) to remove matches less than 500bp in length, and used a second BLAST to identify plasmids with high overall identity: ‘blastn -query ncbi_plasmid.fna -db ncbi_plasmid -perc_identity 93.9 -outfmt 6 -num_threads 8 -out perc_93.9.txt’. We used Python scripts to eliminate matches that did not cover the full length of at least one of the plasmids in this latter BLAST. A Python script compared the results of the two BLAST analyses and removed entries from the first results list if the same plasmid pair appeared in the second results list, generating the final set of mosaic regions. We considered plasmids containing at least one mosaic region mosaic plasmids.

### Gene Detection

All plasmid replication genes and their associated incompatibility groups were identified using PlasmidFinder (16) with the default parameters of 95% nucleotide identity over 60% of the gene length.

Though many transposase sequences have been documented we were unable to identify a transposase sequence database that we could use to confidently identify all transposase sequences in our plasmid sequences. We thus created a custom transposase database based on the J. Craig Venter Institute Genome Propterties (17) database entry for transposase (GenProp1044). All but two elements of this were specific pfams (25) or TIGRFAMs (17). The remaining two profiles (ISPsy8 and IS 150 transposase orfA) had no associated profile HMM. We identified 250 relevant sequences for each from NCBI, created a multiple sequence alignment with the COBALT (26) web interface, and built profile HMMs using HMMER (27).

We downloaded the antibiotic resistance HMM database Resfams (19) from (http://www.dantaslab.org/resfams/). Only the ‘core’ database was used. We indexed all HMM databases for searching using the HMMER command ‘hmmpress <DatabaseName>’, and then performed searches of plasmid proteins against each database using the command ‘hmmscan -o $1.txt --tblout $1_table.txt --max --cpu 16 $1.hmm ncbi_plasmid.faa’ where $1 was the database file basename. The flag “-- cut_tc” was used with the transposase database while “--cut_ga” was used with Resfams, as to increase specificity as indicated by the documentation for Genome Properties and Resfams respectively.

### Inverted repeat detection

We use the EMBOSS (18) program ‘einverted’ to identify inverted repeats in our plasmid sequences with the command ‘einverted -sequence ncbi_plasmid.fna -outfile plasmid_inverted -outseq plasmid_inverted.fasta - maxrepeat 25000’. We used a custom script to align these results with the mosaic fragments we previously identified by BLAST analysis.

### Mosaic fragment clustering

We used a version of the BLAST output file created above, perfect_plasmid_matches.txt, filtered to remove self-alignments and alignments of less than 500 bp, as input for the Python script cluster_from_self_BLAST.py. We used this script to identify how frequently fragments of at least 500 bp occurred in the NCBI plasmid database (including only fragments that occurred in the BLAST output in the analysis, meaning they had to occur at least twice in the plasmid population). Using the Python script extract_seqs.py, we extracted the sequences of the identified fragments and BLASTed those sequences against the NCBI plasmid database FASTA file to double check the occurrences of each fragment.

## Authors’ contributions

MWP and DACB generated hypotheses and designed the study. RT analyzed the antibiotic resistance data. MWP created the custom transposase database, identified mosaic plasmids and mosaic fragments, and correlated those results with the presence of transposases and antibiotic resistance genes. MWP and DACB interpreted the results. DACB and MWP created Figure 1. MWP created the remaining figures.

## Acknowledgements

We would like to thank Mary Lidstrom for providing comments and suggestions during preparation of this manuscript. We would also like to thank the Washington Research Foundation and Mistletoe Foundation for postdoctoral fellowships supporting MWP.

---

**Table 2: Inverted repeat detection results**

**Figure S1:** Boxplot of mosaic and non-mosaic plasmids over plasmid length. Lengths are grouped into bins of 50,000 bp. Y-axis is log scale.

**Figure S2:** Histogram of DNA fragments of at least 500 base pairs over their number of appearances (at 99% nucleotide identity) in the NCBI plasmid database. Mosaic fragments are divided by transposase and antibiotic resistance gene content. Occurrences are grouped into bins of 50. Y-axis is log scale.

**Figure S3:** Distribution of mosaic fragment context diversity. Histogram of DNA fragments of at least 500 base pairs over the number of genera they are found in plasmids from. Y-axis is log scale.

**Table S1:** Mosaic plasmids per genus.

**Table S2:** Genetic context of highly abundant mosaic fragments.

